# Discovery of Synergistic and Antagonistic Drug Combinations against SARS-CoV-2 In Vitro

**DOI:** 10.1101/2020.06.29.178889

**Authors:** Tesia Bobrowski, Lu Chen, Richard T. Eastman, Zina Itkin, Paul Shinn, Catherine Chen, Hui Guo, Wei Zheng, Sam Michael, Anton Simeonov, Matthew D. Hall, Alexey V. Zakharov, Eugene N. Muratov

**Affiliations:** Laboratory for Molecular Modeling, Division of Chemical Biology and Medicinal Chemistry, UNC Eshelman School of Pharmacy, University of North Carolina, Chapel Hill, NC, 27599, USA; National Center for Advancing Translational Sciences (NCATS), 9800 Medical Center Drive, Rockville, Maryland 20850, United States

## Abstract

COVID-19 is undoubtedly the most impactful viral disease of the current century, afflicting millions worldwide. As yet, there is not an approved vaccine, as well as limited options from existing drugs for treating this disease. We hypothesized that combining drugs with independent mechanisms of action could result in synergy against SARS-CoV-2. Using *in silico* approaches, we prioritized 73 combinations of 32 drugs with potential activity against SARS-CoV-2 and then tested them *in vitro*. Overall, we identified 16 synergistic and 8 antagonistic combinations, 4 of which were both synergistic and antagonistic in a dose-dependent manner. Among the 16 synergistic cases, combinations of nitazoxanide with three other compounds (remdesivir, amodiaquine and umifenovir) were the most notable, all exhibiting significant synergy against SARS-CoV-2. The combination of nitazoxanide, an FDA-approved drug, and remdesivir, FDA emergency use authorization for the treatment of COVID-19, demonstrate a strong synergistic interaction. Notably, the combination of remdesivir and hydroxychloroquine demonstrated strong antagonism. Overall, our results emphasize the importance of both drug repurposing and preclinical testing of drug combinations for potential therapeutic use against SARS-CoV-2 infections.

## Introduction

Drug combinations have been used with great success in the past to treat infectious diseases, a notable example being human immunodeficiency virus (HIV).^1^ Drug combinations are particularly useful in treating viral infections due to the fact that they can substantially lower the risk of the development of resistance to any one drug, and have demonstrated marked success against various viral diseases in the past.^1,2^ Additionally, the antiviral action of the drug combination may be stronger than either drug alone, a phenomenon known as synergy. Antiviral synergy has been previously illustrated for the treatment of hepatitis C virus (HCV),^2–4^ HIV,^5^ herpes simplex virus (HSV),^6,7^ poliovirus,^8^ Ebola virus,^9^ Zika virus,^10^ and human cytomegalovirus (HCMV).^11^ Often the rationale for why such synergism occurs remains unclear – it is extraordinarily difficult to provide explanations for existing synergistic or antagonistic drug combinations without prior, extensive, experimental investigations.^12^

For antiviral drug synergy predictions, some past evidence suggests that combinations of antiviral drugs that are of different classes, have varied mechanisms of action, and act upon different stages of the virus life cycle, are more likely to be synergistic.^2,3,13^ Though there have been many compounds suggested against SARS-CoV-2 tested *in vitro*, many of these drugs are only being evaluated as single agents, according to ClinicalTrials.gov. These include numerous research groups evaluating compounds in enzymatic and cellular assays to determine antiviral activity.^14–23^ Comparatively, there has been limited systematic screening of drug combinations.^24^

Meanwhile, the situation in clinical trials is somewhat different. Of the 2,341 clinical trials relevant to COVID-19 as of June 27, 2020, ca. 100 describe drug combinations. However, many of these trials are evaluating the same drug repurposing combinations, e.g., lopinavir+ritonavir or azithromycin+hydroxychloroquine.^25^ Perhaps the most noteworthy antiviral drug combination to date has been lopinavir+ritonavir (formulated as a single therapeutic, Kaletra), which has been tested in clinical trials with and without Interferon-β1b.^26^ This Phase II trial for a triple antiviral therapy combining interferon-β1b, lopinavir–ritonavir, and ribavirin was shown to shorten the duration of viral shedding and hospital stay in patients with mild to moderate COVID-19.^26^ However, a recent study assessing the effectiveness of the lopinavir+ritonavir combination in treating COVID-19 has noted that administration of lopinavir+ritonavir in COVID-19 patients showed no benefit.^27^ As of June 27, 2020, no combination therapy has yet yielded positive results in Phase III randomized clinical trials.^28^

While there are many ongoing or upcoming clinical trials testing combinations to treat COVID-19, few have undergone extensive preclinical studies prior to their combination in patients. Due to a lack of such studies, more information is needed on the combinatorial use of antivirals and other drugs against SARS-CoV-2 in order to (1) more efficiently prioritize synergistic combinations for translation into clinical use; and (2) flag antagonistic combinations prior to their evaluation in the preclinical stage. To this point, we have recently used data and text mining approaches to propose drug combinations for repurposing against SARS-CoV-2,^29^ operating on the assumption that combinations of drugs with differing mechanisms might exhibit synergistic activity. The goal of this study is to report the antiviral activity, synergy, and antagonism of 73 binary combinations of 31 drugs identified earlier^29^ as demonstrated in an *in vitro* SARS-CoV-2 cytopathic assay.

## Methods

### Selection of drug combinations

We applied a combination of text mining (Chemotext),^30^ knowledge mining (ROBOKOP knowledge graphs),^31^ and machine learning (QSAR)^32^ in the search for existing drugs with possible activities against SARS-CoV-2. A detailed description of our study design is provided in our recent paper and is also depicted in Fig. 1.^29^ Briefly, we first identified a list of 76 individual drug candidates for repurposing in combination therapy against COVID-19 using aforementioned techniques. Here we selected some of the hits from virtual screening of DrugBank^33^ and NPC^34^ collections by our QSAR models of SARS-CoV M^pro^ inhibition^35^ and all the drugs found in Chemotext and ROBOKOP searching for “SARS”, “*Coronaviridae”*, etc. This resulted in 76 individual drugs that may potentially create 2580 binary and 70300 ternary combinations. In this study we will describe only the binary combinations.

**Figure 1.**
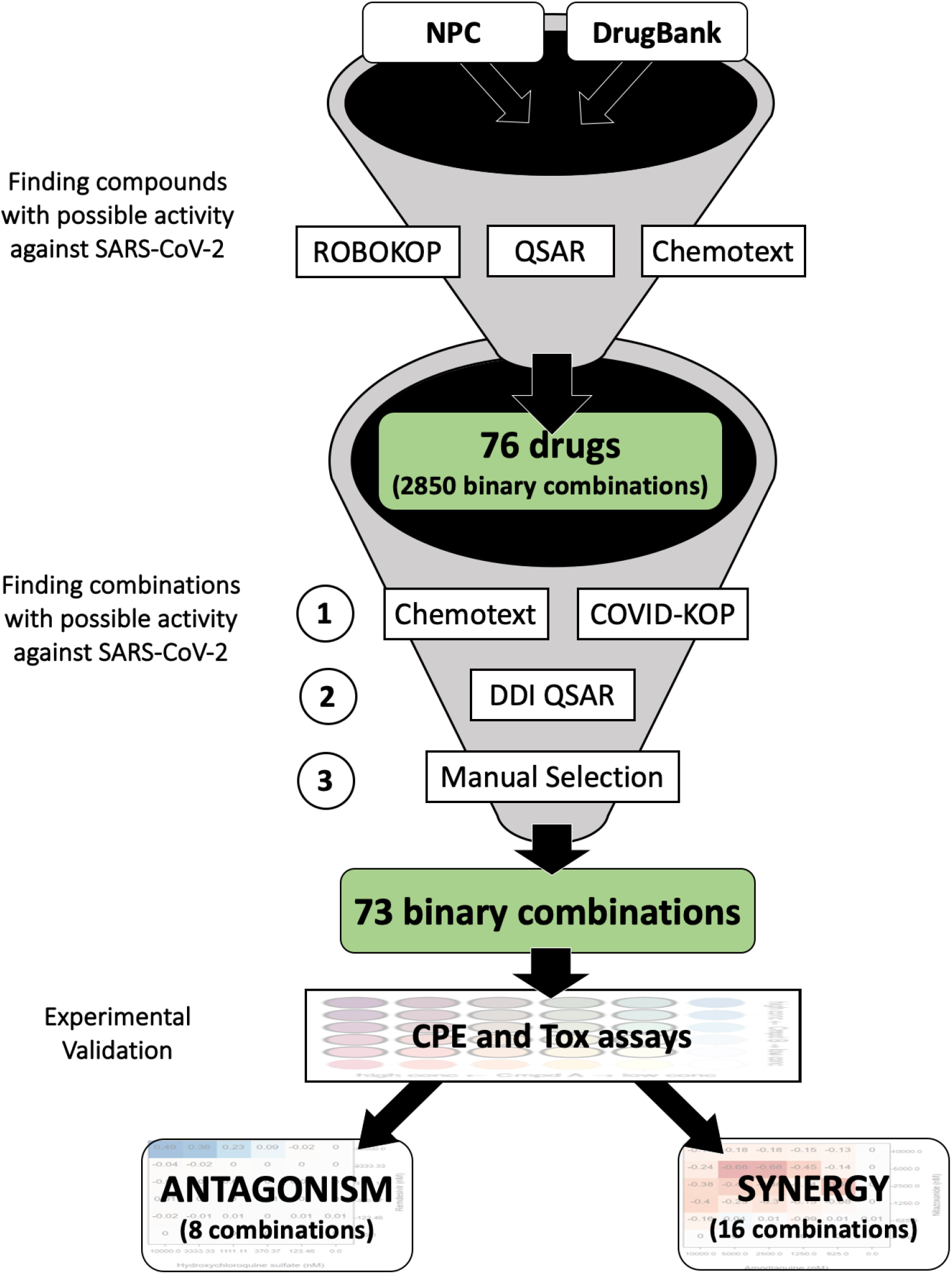
Study design for selecting possible synergistic drug combinations. In this study we report only 73 binary combinations. 95 ternary combinations identified in a similar fashion will be reported in a future study.

Then, we applied Chemotext and ROBOKOP to evaluate the potential combinations of selected drugs. We also applied the recently developed web-platform COVID-KOP^36^ to refine the original list of combinations. Here, we considered the mechanism(s) of action (if known) and the target(s) of the drugs (a recent review^37^ was extremely helpful here) to identify potential synergistic effects and avoid antagonistic interactions.^24,38^ We aimed (whenever possible) to prioritize combinations of drugs with different mechanisms of action and/or targeting the virus at different stages of its lifecycle (higher confidence hits) or at least acting upon distinct viral protein targets, which increases the probability of synergy between drugs.^39,40^ This rationale behind combination selection is depicted in Fig. 2 through the example of umifenovir and emetine, which are suspected to act upon different stages of the viral lifecycle (see Fig. 2).

**Figure 2.**
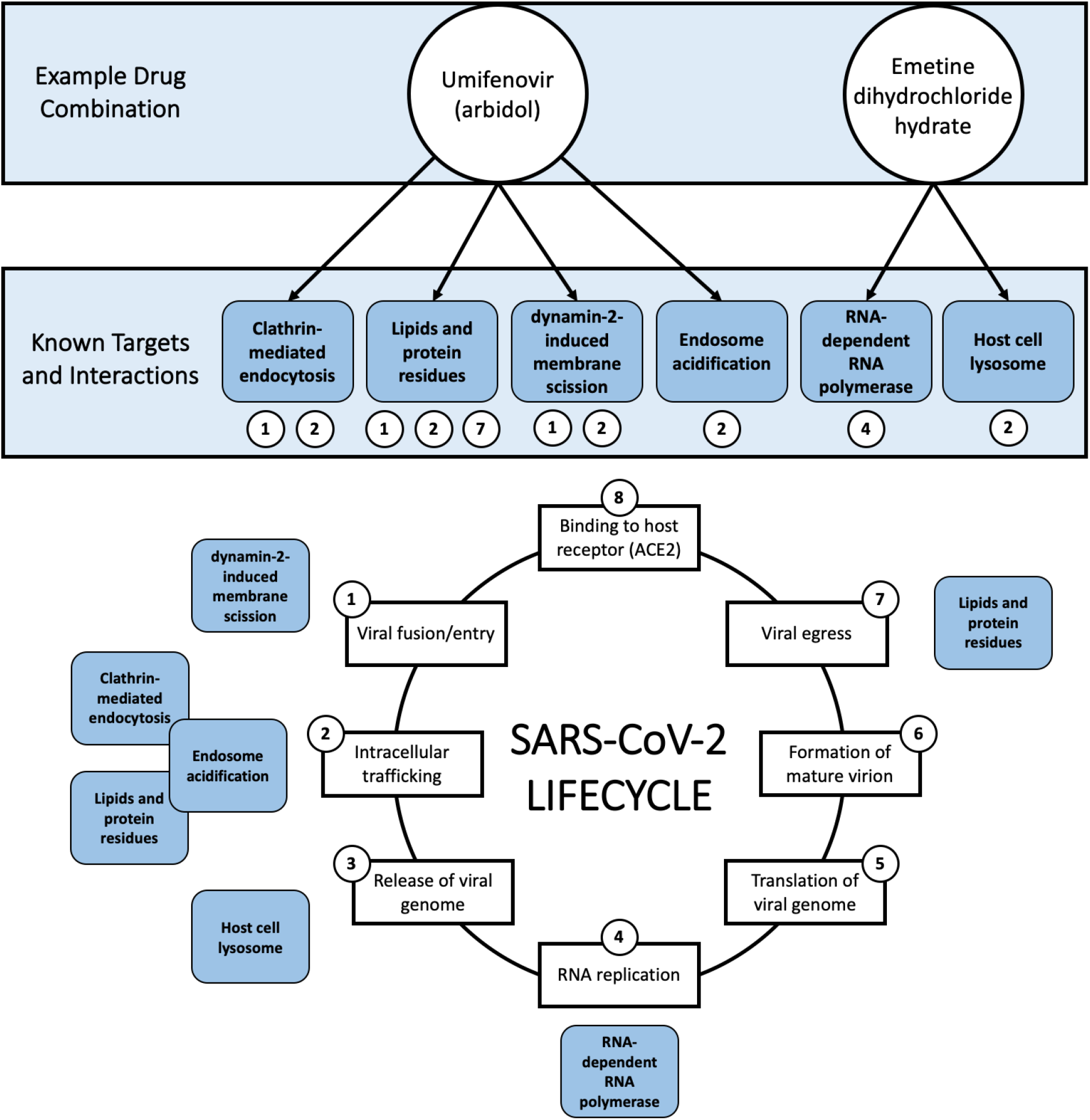
Example (umifenovir + emetine dihydrochloride hydrate) of the rationale behind mixture selection, i.e., interference with different steps of the COVID-19 lifecycle. All known targets and interactions were taken from the literature and are not necessarily specific to SARS-CoV-2. Umifenovir’s proposed mechanisms of action involve clathrin-mediated endocytosis,^41,42^ lipids and protein residues,^43^ dynamin-2-induced membrane scission,^41^ and endosome acidification.^41^ Emetine’s proposed mechanisms of action involve RNA replication/RNA-dependent RNA polymerase^44,45^ and the host cell lysosome.^45^ The viral lifecycle of SARS-CoV-2 was inspired by Fig. 1. of da Costa et. al 2020.^46^

Next, we used Chemotext to determine if these compounds had been tested together before. One capability of Chemotext is finding papers that share two search terms, in this case, the names of compound #1 and compound #2, and returning these common papers by the MeSH terms found in these papers. These shared MeSH terms, depicting proteins, chemicals, etc. further allow us to hypothesize on how these two compounds may be connected, namely via shared biochemical pathways. Cheminformatics models were then used to further exclude combinations with undesirable drug-drug interactions.^47,48^ Subsequently, combinations were manually curated to exclude artifacts as well as compounds undesired or contraindicated in the case of pneumonia (e.g., paclitaxel, bleomycin), or viral infection (e.g., baricitinib). In the end, we prioritized 73 binary combinations of 32 drugs for further experimental testing. Computational approaches used for prioritization of combinations and mechanistic explanations are briefly outlined below.

**Chemotext** is a publicly-available web server that mines the published literature in PubMed in the form of Medical Subject Headings (MeSH) terms.^30^ Chemotext was used for elucidation of the relationships between drugs and their combinations, targets, and SARS-CoV-2 and COVID-19 from the papers annotated Medline/PubMed database.

**ROBOKOP**^49^ and **COVID-KOP**^36^. ROBOKOP is a data-mining tool developed within Biomedical Data Translator Initiative^50^ to efficiently query, store, rank and explore sub-graphs of a complex knowledge graph (KG) for hypothesis generation and scoring. We have used ROBOKOP in a similar fashion as Chemotext; Chemotext could help the user to find and impute the connections between drugs, targets, and diseases and ROBOKOP could help explore and score them. In the middle of the project, COVID-KOP, a new knowledgebase integrating the existing ROBOKOP biomedical knowledge graph with information from recent biomedical literature on COVID-19 annotated in the CORD-19 collection,^51^ was developed and thus we began to utilize it instead of ROBOKOP.

**QSAR models** developed by us earlier were used for selection of drugs^35,48^ that could be repurposed as combinations and exclusion of potential drug-drug interactions and side effects.^52^ All the models were developed according to the best practices of QSAR modeling^32,53,54^ with a special attention paid for data curation^55–57^ and rigorous external validation.^58^ Mixture-specific descriptors and validation techniques^59^ specially developed for modeling of drug combinations were utilized for modeling of drug-drug interactions.^52^

### Assay-ready plate production

An in-house software package, Matrix Script Plate Generator (MSPG), was used to create all necessary files for generation of compound source plates, acoustic dispensing scripts, and plate maps for each combination in 6 × 6 dose matrix, shown in Fig. 3H. Briefly, we placed single-agent compounds A and B (1:3 or 1:2 dilution) in orthogonal directions, and used the remaining 25 wells to measure the combinatorial outcome of A+B with their respective doses.

**Figure 3.**
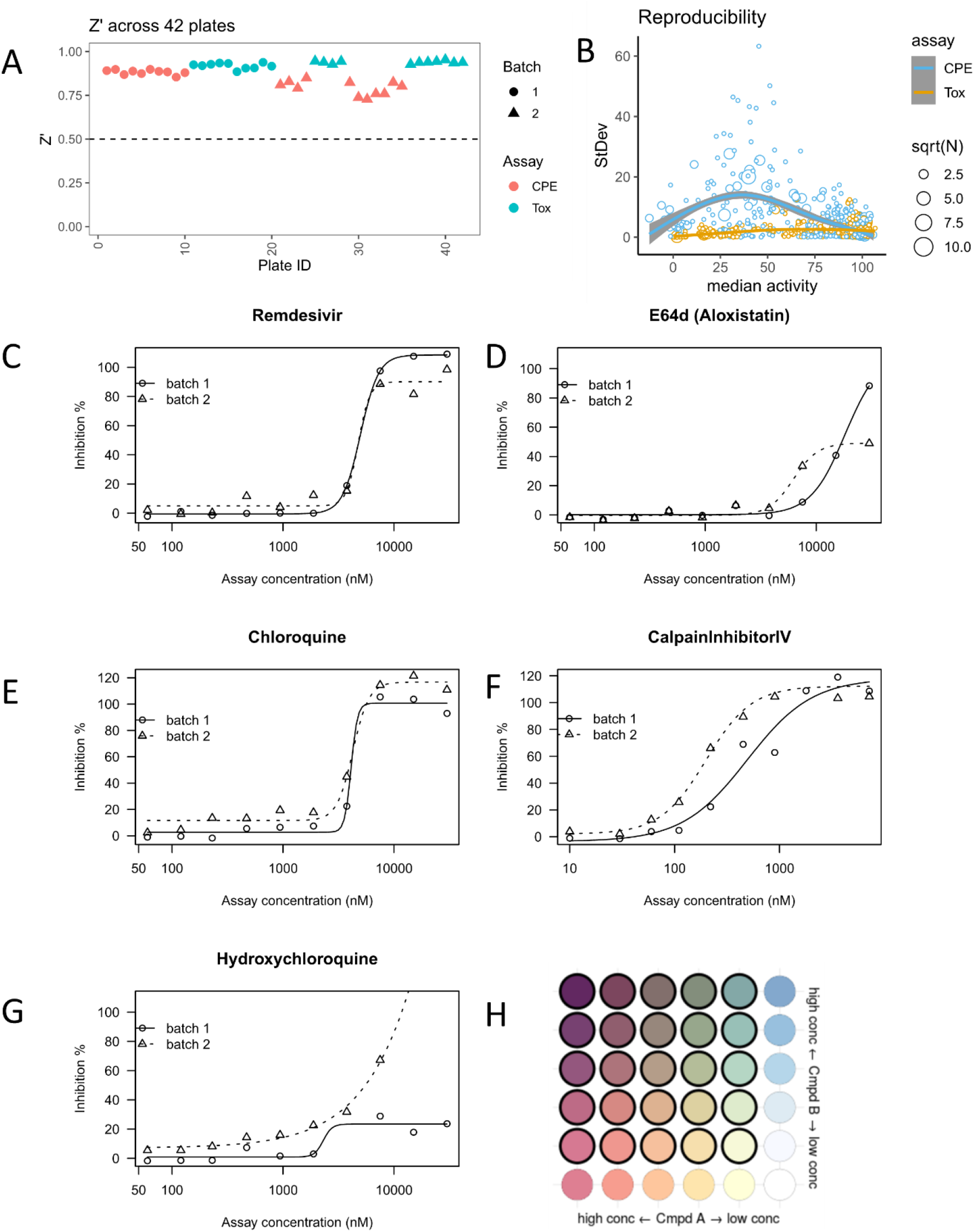
Performance of matrix screening. (A) Z’ factor on different assays (CPE or Tox) and biological batches (B) Reproducibility across all replicates (defined as a compound at certain concentration). Number of replicates (N) may vary, e.g., more single-agent replicates were performed due to matrix setting. (C-G) Dose response curves from an independent benchmark set performed at a different site. (H) layout of a 6 × 6 dose matrix. Wells with (or without) bold border represent dose combination (or single agent alone).

To generate the compound source plate, a Perkin-Elmer Janus Automated Workstation was used to transfer compounds from 1.4 mL Matrix 2D barcode tube (sample tube) to the individual wells of Echo Qualified 384-Well Polypropylene Microplate 2.0**, along with positive and neutral (DMSO) controls (Labcyte, San Jose, CA). The plates were briefly centrifuged for 2 min at 100×g. Customized generated input files for the Labcyte acoustic dispenser (Labcyte, San Jose, CA) from a comma-separated value file that contained the unique plate and well pairings for each of the initial compound matrix blocks. Compounds were dispensed to generate assay ready plates using an Access Laboratory Workstation with dual Echo 655 dispensers (Labcyte). Each plate was sealed with a peel-able aluminium seal, remaining covered until the initiation of the biological assay, frozen at −80°C until used for screening.

### Viral <u>C</u>ytopathic <u>E</u>ffect assay (CPE) and cytotoxicity assay (Tox)

The detailed protocol for the CPE and Tox counter assays are available from NCATS OpenData portal.^23^

CPE: https://opendata.ncats.nih.gov/covid19/assay?aid=14

Tox: https://opendata.ncats.nih.gov/covid19/assay?aid=15

Briefly, 30 nL of each compound in DMSO was acoustically dispensed into assay ready plates (ARPs). Media was then added to the plates at 5 μL/well and incubated at room temperature to dissolve the compounds. Vero E6 cells (selected for high ACE2 expression) were premixed with SARS-CoV-2 (USA_WA1/2020) at a MOI of 0.002, and were dispensed as 25 mL/well into ARPs within 5 min in a BSL-3 lab. The final cell density was 4000 cells/well. The cells and virus were incubated with compounds for 72 hrs, then viability was assayed by ATP content.

For the cytotoxicity assay, ARPs were prepared in the same way as for CPE assay. Then, 5 mL/well of media was dispensed into assay plates. Vero E6 in media and dispensed into assay plates at 25 μL/well for a final cell density of 4000 cells/well. Assay plates were incubated for 72 hrs at 37 °C, 5% CO_2_, 90% humidity, before viability was assayed by ATP content.

### Data analysis

CPE and Tox activity were normalized using independent control wells on each plate, so activity values were not strictly bounded between [0, 100]. For CPE assay, DMSO+virus was treated as the neutral control, whereas DMSO-only (no virus) served as the positive control. A Calpain inhibitor IV was used as batch control (2ug/ml final assay concentration). Normalized CPE activity = 1 − (x − neutralCtrl) / (positiveCtrl − neutralCtrl) × 100%. For Tox assay, DMSO-only was used as the neutral control and media-only wells (no cell) as the negative control. Normalized Tox activity = (x − negativeCtrl) / (neutralCtrl − negativeCtrl) × 100%. Plate-level data was pivoted to block-level data and replicates were median-aggregated.

Synergism and antagonism from a 6 × 6 block were evaluated using the highest single agent model (HSA).^60^ Given a dose combination A_conc1_ + B_conc2_,

HSA(A_conc1_+B_conc2_) = activity(A_conc1_+B_conc2_) – MIN{activity(A_conc1_), activity(B_conc2_)}

Synergism: HSA(*) < 0

Antagonism: HSA(*) > 0

Additivity: HSA(*) = 0

To account for dose-dependent synergism and antagonism, we analyzed the negative HSA (HSA.neg) and positive HSA (HSA.pos) separately. The overall synergism (or antagonism) given a 6 × 6 block was calculated as the sum of all negative (or positive) HSA(A_conc*_+B_conc*_) across the non-toxic dose combinations (defined as Tox activity > 50).

Since CPE activity showed a non-linear variation across different activity levels (Fig. 3B), which made it more difficult to ascertain the reproducibility of synergy/antagonism (given limited resources), we evaluated the smoothness of the 2D activity landscape. Smoothness was calculated as the root mean squared deviation (RMSD) between the actual and gaussian smoothed (σ = 2) landscape. If a block had RMSD(observed, gaussian smoothed) > 20 or less than 25 non-toxic CPE values, inconclusive synergism/antagonism was recorded. Otherwise, synergism and/or antagonism were recorded if HSA.neg < −100 and/or HSA.pos > 100.

## Results

### Performance of matrix screening

In total, we screened 73 pairwise combinations in a 6×6 dose matrix format, which involved two biological batches (cell and SARS-CoV-2 virus) and two assays (cytopathic effect and cytotoxicity against Vero-E6 cells) across 42 384-well plates including replicates (Supplementary File 1 and Supplementary File 4). The Z’ factor was robust across batches and assays (all Z’ > 0.7, Fig. 3A). Each batch was assessed by a benchmark compound collection including five known antivirals, performed at an independent site. We did not observe significant drift of potencies or efficacies between batches, except for hydroxychloroquine, which consistently resulted in inconclusive dose response curves (Fig. 3C-G and Supplementary File 2). In addition, we performed a third QC to check the reproducibility across all available replicates in CPE or Tox assays (Fig. 3C). CPE activity showed a biphasic trend between median activity and standard deviation: most reproducible (StDev < 20) when activity is extreme (activity < 20 or > 75), and less reproducible in between (blue points). This is probably due to the high sensitivity to technical/biological variations when the concentration is close to the EC_50_. This biphasic reproducibility of CPE activity also highlights the importance of using a dose matrix, instead of a single dose combination, to enhance the confidence of synergism/antagonism findings. In contrast, Tox activity is highly reproducible regardless of the median activity (yellow points).

### Overview of hits

Since synergism and antagonism might occur simultaneously in a concentration-dependent manner (see Fig. 7 for an example), we separated the synergism and antagonism analyses using highest single agent (HSA) synergy model. Within 73 binary combinations of 32 compounds, we identified 16 synergistic and 8 antagonistic combinations, 4 of which displayed both synergistic and antagonistic interactions at different compound concentrations (Fig. 4 & 5). There are 29 combinations with inconclusive determinations that require additional validation due to cytotoxicity or the roughness of the activity landscape (see Methods). A summary of the screening result is available in Supplementary File 4. A map of drug combinations depicting their synergism/antagonism outcomes is depicted in Fig. 5.

**Figure 4.**
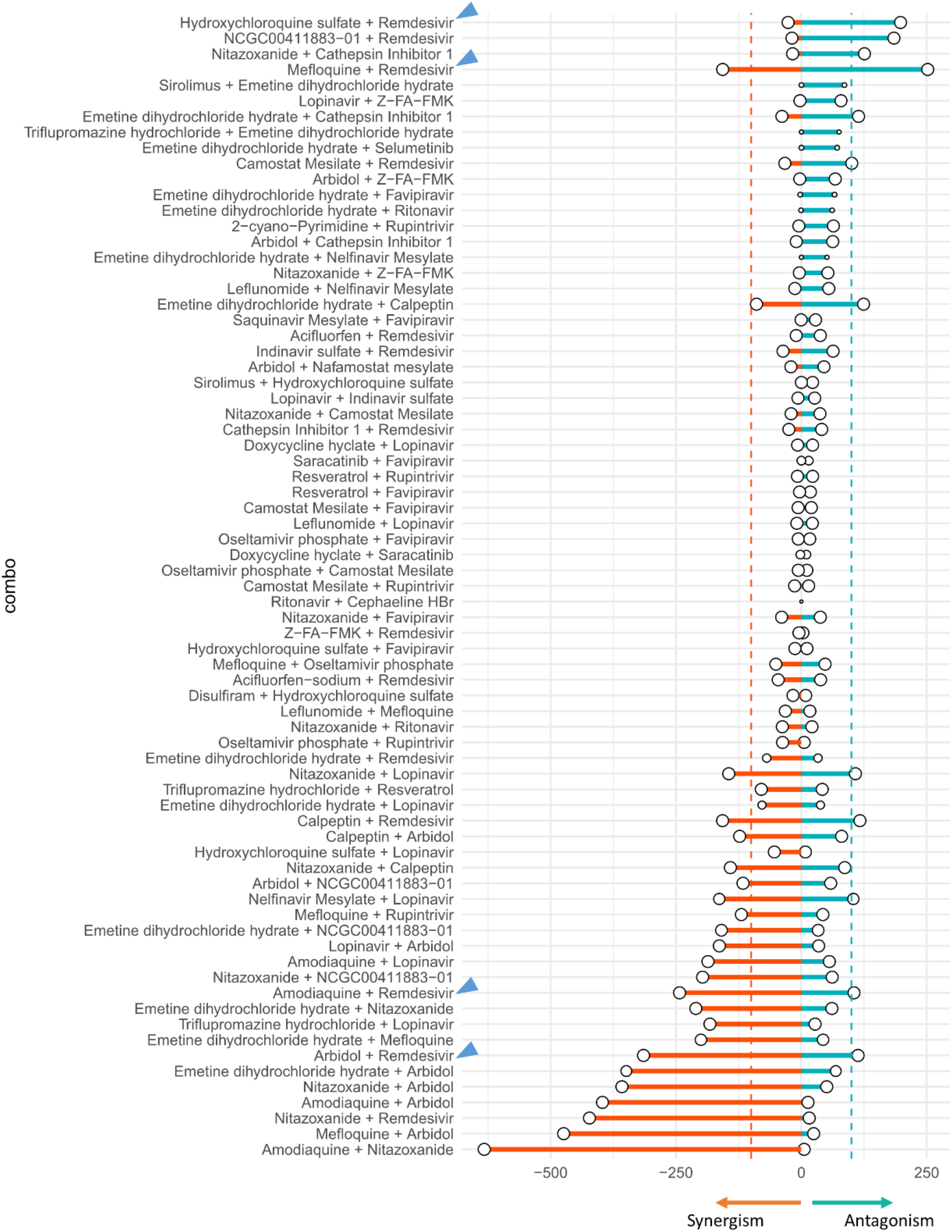
Summary of synergism or antagonism across 73 tested combinations. Due to biphasic dose response, synergism was separated from antagonism. Synergism is calculated as the sum of HSA.neg values from non-toxic dose combinations (Tox > 50%), and vice versa. The size of circle reflected the confidence of the observed synergism/antagonism (bigger circle = less doses were excluded due to toxicity).The inconclusive blocks (N_nontoxic_ < 25 or rough activity landscape) were shaded. Two dashed lines indicated the cutoff of HSA synergism (−100) or antagonism (100). Blue arrows highlighted the combinations between remdesivir and tertiary amine compounds from conclusive blocks.

**Figure 5.**
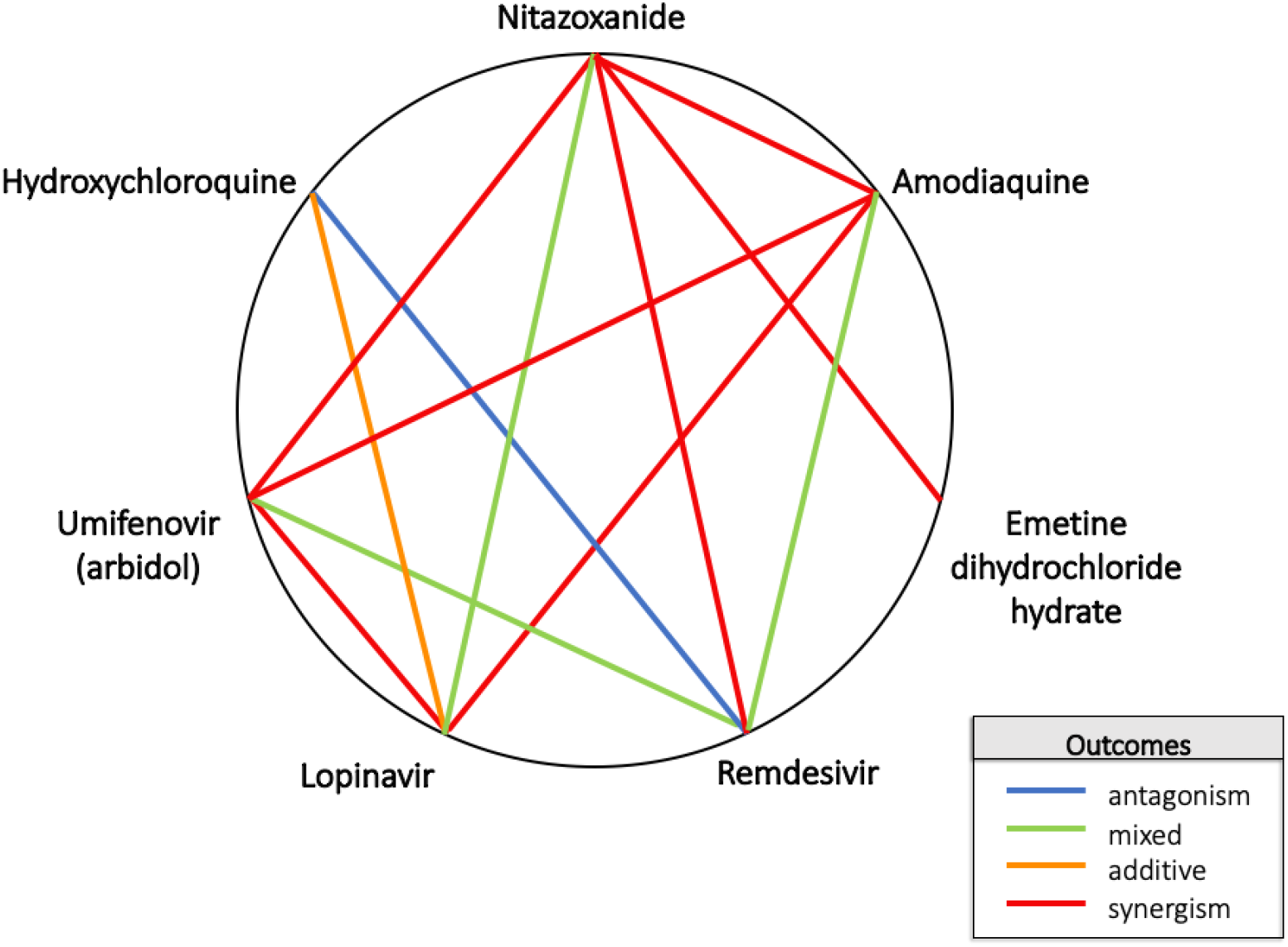
Heptagonal polygonogram depicting some of the binary combinations tested in the study. Degrees of synergy/antagonism were ascertained from Figure 4. The definitions were defined based on the degree of HSA synergism/antagonism determined in the CPE assay.

Next, we investigated whether any drug combinations with different mechanisms of action (MoA) could offer a greater chance of antiviral synergism or antagonism. A previous study demonstrated that synergism/antagonism is predictable based on MoA in oncology screening.^61^ Unfortunately, we found limited evidence of MoA-associated synergism/antagonism (up to 10 μM) for SARS-CoV-2 (Fig. 6). The most antagonistic MoA combination came from the combination of an RNA-dependent RNA polymerase (RdRp) inhibitor (remdesivir) and an antimalarial drug, in which 3 (hydroxychloroquine, amodiaquine and mefloquine) out of 4 appeared to be antagonistic (Fig. 7). However, the current data are not sufficient to conclusively infer any MoA-associated synergism/antagonism. We are performing further systematic synergy screening to evaluate combinations with known mechanisms of action.

**Figure 6.**
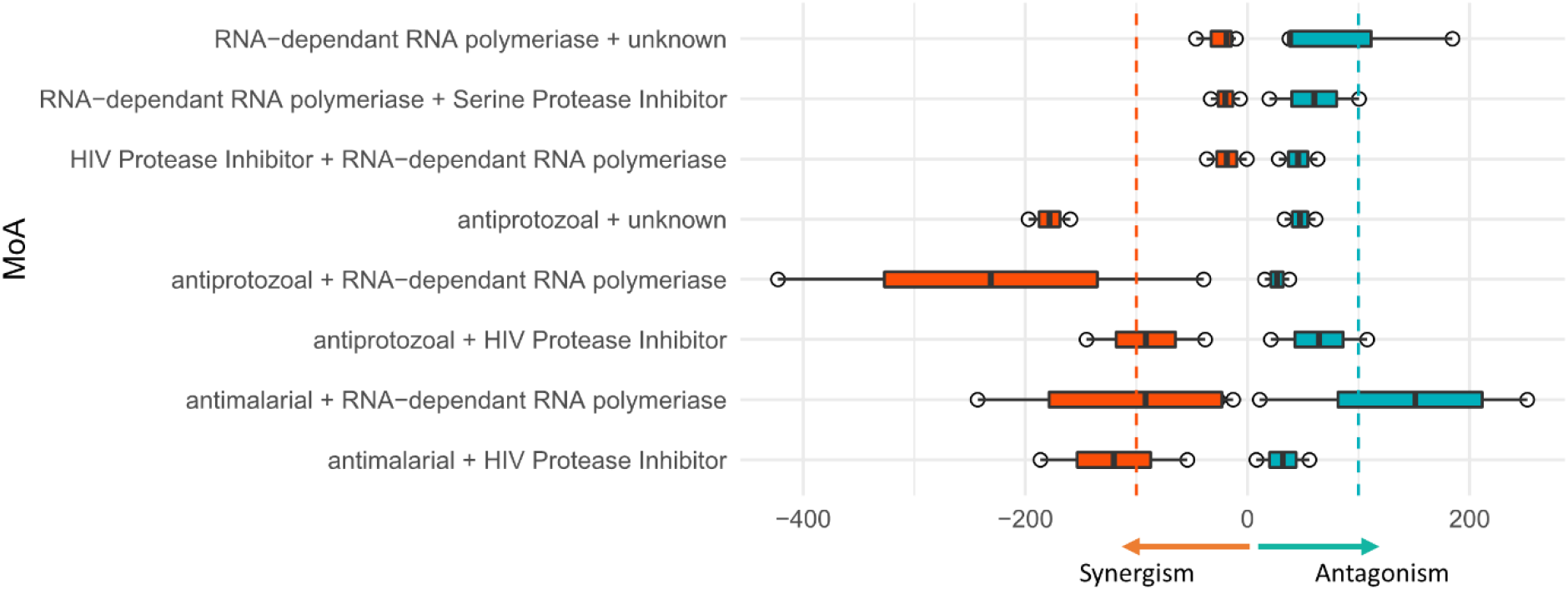
Summary of synergism or antagonism over different mechanism of action (MoA) combination. Inconclusive blocks or singleton MoA was excluded. Two dashed lines indicated the cutoff of HSA synergism (−100) or antagonism (100).

**Figure 7.**
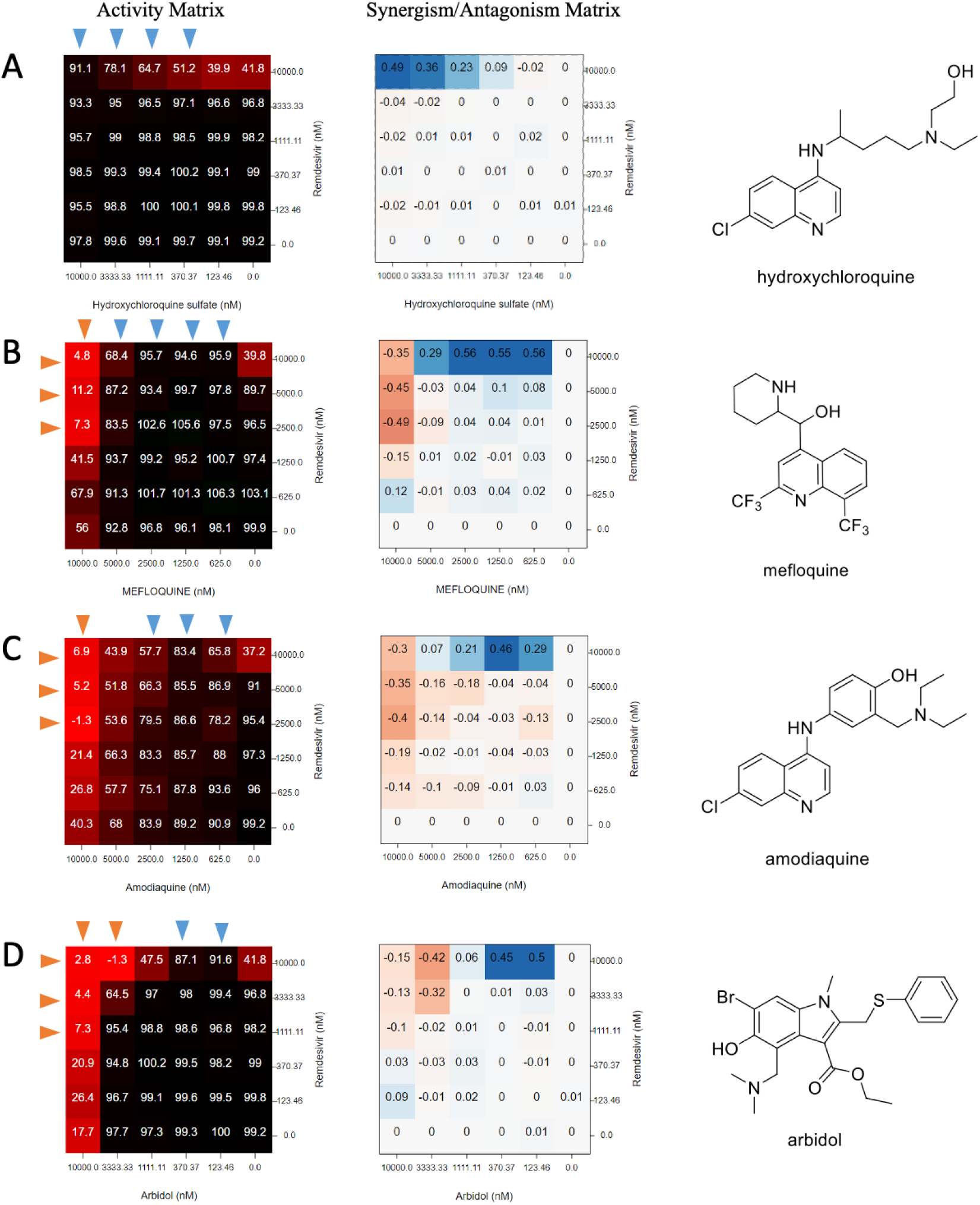
Matrix blocks from Remdesivir + amine drugs in CPE assay. The activity was normalized so that 100: virus fully viable and 0: no virus. Red arrow: the concentrations that synergize with the partner compound. Blue arrow: the concentrations that antagonize against the partner compound. Chemical structures were shown on the right. (A) hydroxychloroquine (B) mefloquine (C) amodiaquine (D) arbidol.

### Antagonism between remdesivir and lysosomotropic agents

Most notably, our results demonstrate a strong antagonistic effect between remdesivir and antimalarial drugs, including hydroxychloroquine, mefloquine, and amodiaquine (Fig. 7). The most striking antagonism was observed in the combination of the only two drugs ever approved with FDA Emergency Use Authorization (EUA): hydroxychloroquine and remdesivir (though we note the EUA for hydroxychloroquine has been withdrawn by the FDA).^62^ Our results showed that 10 μM hydroxychloroquine could completely extinguish the antiviral activity of remdesivir in vitro. The antagonistic effect exerted by hydroxychloroquine could be observed at a concentration as low as 0.37 μM (Fig. 7A). In contrast to hydroxychloroquine, which only showed antagonism under 10 μM, mefloquine and amodiaquine could synergize with remdesivir at high concentrations (Fig. 7B and Fig. 7C). This biphasic interaction pattern (antagonism at low concentration and synergism at high concentration) ruled out the possibility of direct chemical interaction between remdesivir and tertiary amines in a mixture.

COVID-KOP was further utilized to seek possible explanations for the observed synergies and antagonisms in the study. A pertinent use of COVID-KOP is to identify the biological processes and activities common to two drugs, which could suggest possible common interactions that lead to antagonism, such as in the case of remdesivir and hydroxychloroquine, shown in Fig. 8. Analysis using COVID-KOP showed that hydroxychloroquine and remdesivir are both associated with the biological process/activity terms: “clathrin-dependent endocytosis”; “viral entry/release into host cell”; “inflammatory response”; “pH reduction”; “negative regulation of kinase activity”; and “protein tyrosine kinase activity”, as well as different interleukin receptor (1-2, 6-7, 10) binding events (see Fig. 8). This computational analysis indicates that these are common terms to both drugs, meaning that one or more of these biological processes are possible points where these two drugs could interact to cause antagonism. This type of network analysis provides bird’s-eye view of all the possible pathways or processes implicated in outcomes such as antagonism and synergism.^63^ These terms cannot be resolved further; thus, it may be possible that the action of a protein kinase is involved in the antagonism observed for the combination of remdesivir and hydroxychloroquine (this is considered further in the Discussion), but it is unclear which specific protein kinases this entails based solely on COVID-KOP results. It should be noted that with more widely known compounds, Chemotext, ROBOKOP, and COVID-KOP are more likely to return shared terms that may not have real significance. That is, given that remdesivir has been mentioned in many COVID-19 studies that may not necessarily focus on the drug’s mechanism of action, the connections it has to certain targets may be somewhat spurious. This is also true for hydroxychloroquine, which was also widely touted as a possible treatment early in the pandemic.

**Figure 8.**
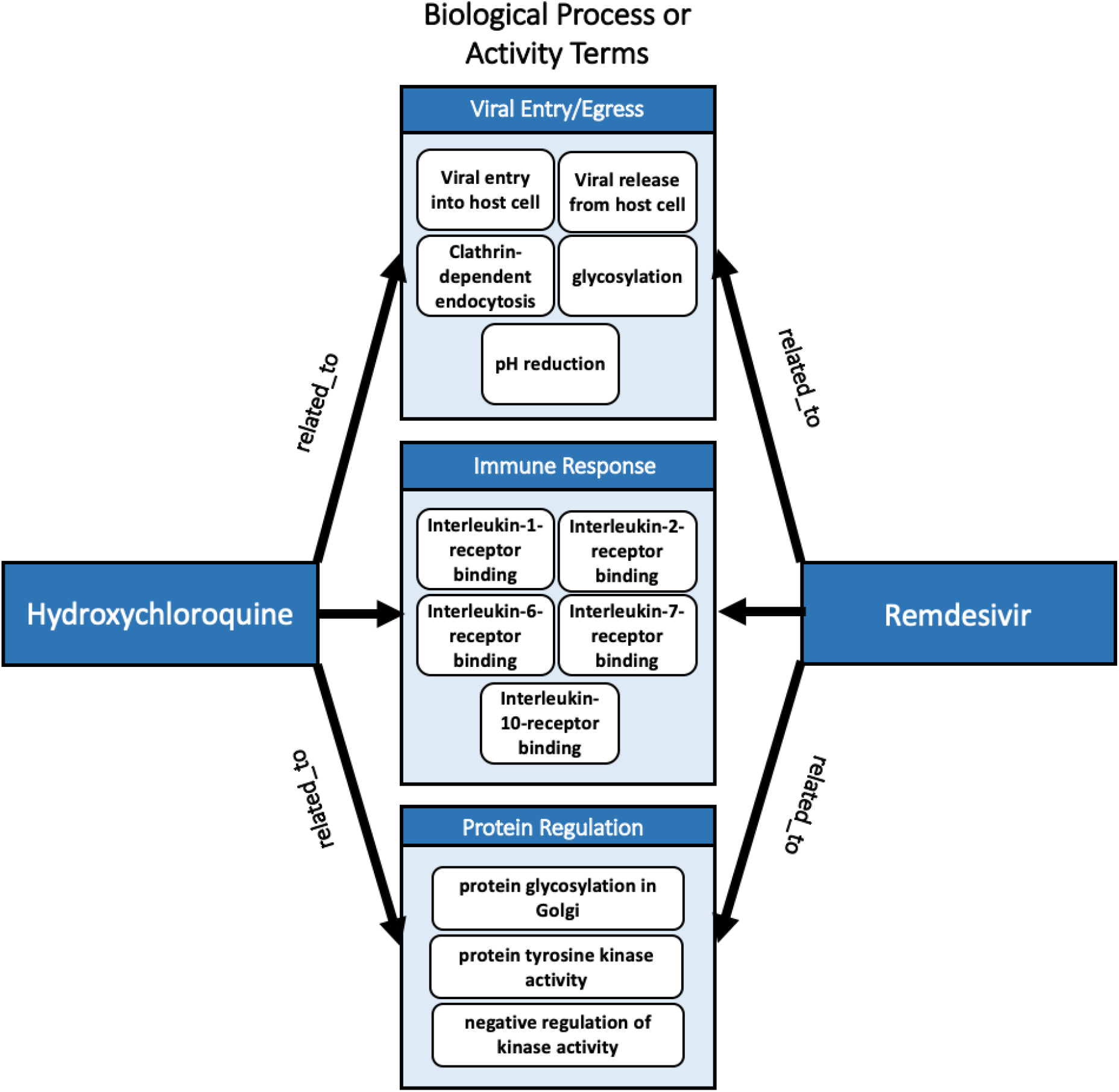
COVID-KOP identification of “biological process or activity” terms related to hydroxychloroquine and remdesivir. Relevant results are displayed and categorized by type. Common terms indicate processes/activities in which hydroxychloroquine and remdesivir may interfere with each other’s’ mechanisms, resulting in antagonism.

We observed a similar antagonism against remdesivir from another drug with a tertiary amine moiety, umifenovir (arbidol, antiviral approved in Russia/China), at low concentrations (123 - 370 nM, Fig. 7D). However, umifenovir synergized with remdesivir at high concentrations (3 - 10 μM). Hydroxychloroquine, mefloquine and amodiaquine are known lysosomotropic agents^64,65^ and umifenovir also contains a tertiary amine moiety, suggesting an association between reduced antiviral efficacy of remdesivir by lysosomotropic amines.

### Synergism between nitazoxanide and remdesivir, amodiaquine, or umifenovir

Among 16 synergistic combinations, we observed the strongest synergistic effects in the combinations containing nitazoxanide, and FDA-approved broad-spectrum antiviral and antiparasitic drug. The three most synergistic combinations with nitazoxanide, i.e., nitazoxanide + remdesivir / umifenovir / amodiaquine are shown in Fig. 9. A complete rescue of CPE could be observed from 0.625 - 5 μM of nitazoxanide when combined with remdesivir / umifenovir / amodiaquine, where any of these drugs alone could only achieve a maximum 40-60% rescue (Fig. 9A-C). Nitazoxanide is not cytotoxic when concentration is below 5 μM. However, we observed a mild toxicity (~20%) of nitazoxanide at 10 μM (Supplementary File 3), which may explain the vanishment of synergy at this concentration.

**Figure 9.**
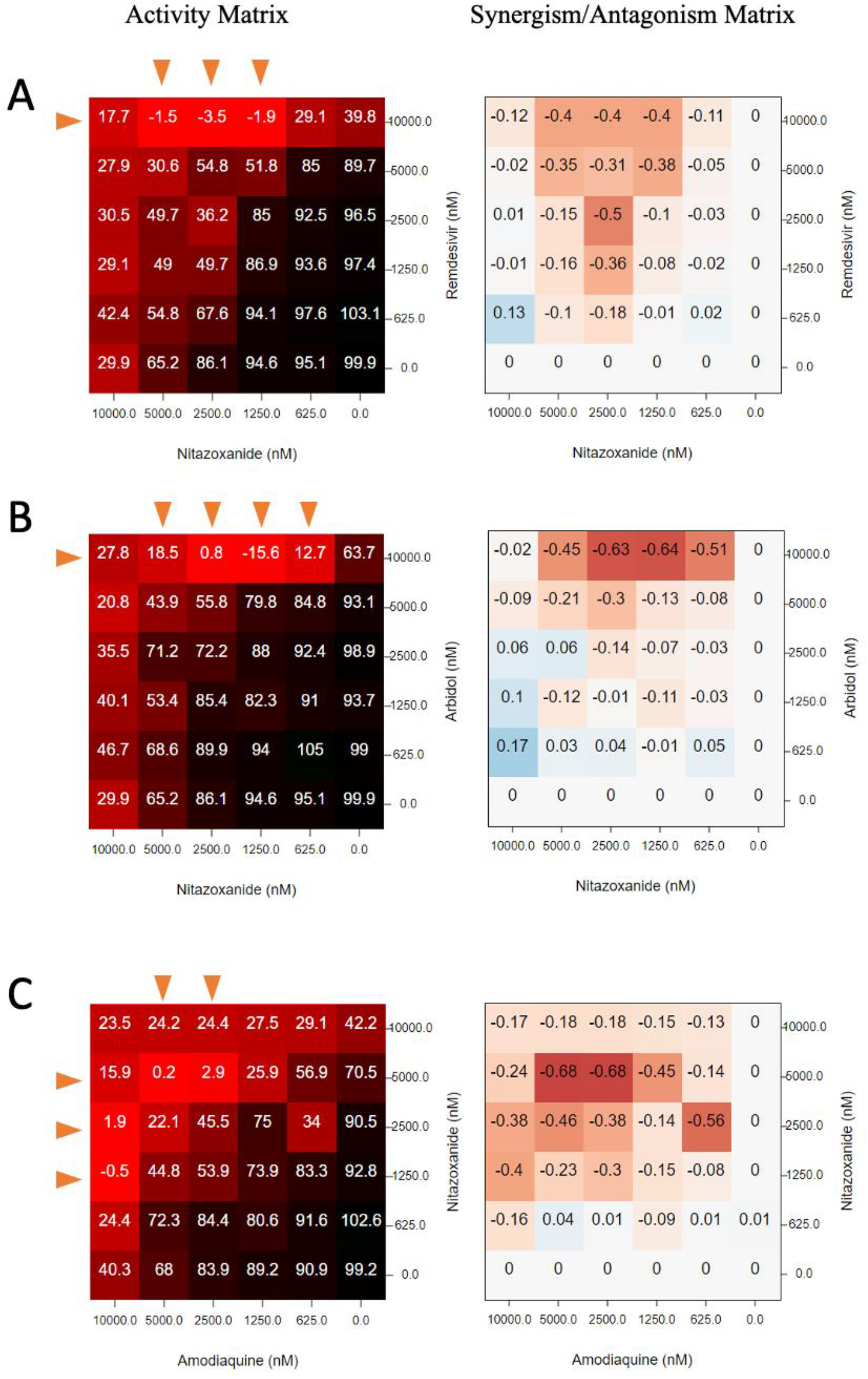
Matrix blocks from 3 synergistic combination involving nitazoxanide. (A) nitazoxanide + remdesivir; (B) nitazoxanide + arbidol; (C) nitazoxanide + amodiaquine. Red arrow: the concentrations that synergize with the partner compound.

## Discussion

Combining modern computational techniques and experimental approaches, we have identified sixteen synergistic antiviral combinations (see Fig. 4). Somewhat unexpectedly, our results also revealed an antagonism between remdesivir and hydroxychloroquine, the two drugs approved with FDA Emergency Use Authorization for treatment of COVID-19 (the EUA for hydroxychloroquine has since been rejected, as of June 15, 2020^62^). Remdesivir also exhibits antagonism with lysosomotropic agents and some other drugs. These findings demonstrate the importance of preclinical research investigating antiviral drug combinations prior to their application in patients, as well as the utility of data and text mining approaches to explore MoAs underlying synergism/antagonism within the context of COVID-19. The approach offered here indicates that when seeking synergistic antiviral drug combination therapies, it is useful to combine drugs that act upon different parts of the viral lifecycle.^66^ These data also emphasize the utility of drug repurposing in treating COVID-19, but also its pitfalls; namely, while individual drugs may be safe for use in patients, combinations of these drugs may not exhibit the same safety or efficacy profile. Lack of preclinical studies on combinations prior to their administration in patients may significantly increase the risk of antagonism and undesirable side effects. The matrix screening platform presented in this study is an efficient, data-driven method for prioritizing synergistic combinations and flagging undesirable drug-drug interactions.

For instance, our method was successful in identifying the antagonistic effect of remdesivir and hydroxychloroquine in combination. Remdesivir is a nucleotide analogue prodrug, which inhibits SARS-CoV-2 RNA-dependent RNA polymerase (RdRp) through inducing delayed chain termination.^67^ Although the key enzymes required to activate remdesivir into its active triphosphate form have not been reported, we hypothesize that remdesivir shares, at least partially, a similar activation pathway with GS-465124, a metabolite from nucleotide prodrug GS-6620 for hepatitis C virus, based on the chemical similarity (Fig. 10).^68^ Both remdesivir and GS-465124 are adenosine-like phosphoramidates, with the only difference being a methyl group on the 2’ pentose ring. Therefore, remdesivir is also likely to be hydrolyzed into alanine metabolite (GS-704277) intracellularly by cathepsin A, akin to GS-465124.^68^ Since cathepsin A is an acidic pH-dependent serine protease strictly located in lysosome, the lysosomotropic agents such as hydroxychloroquine may reduce the amount of active metabolite by increasing lysosomal pH.^69^ More mechanistic studies of remdesivir are in progress at NCATS, utilizing label-free mass spectra to elucidate the exact mechanisms underlying this striking antagonism.

**Figure 10.**
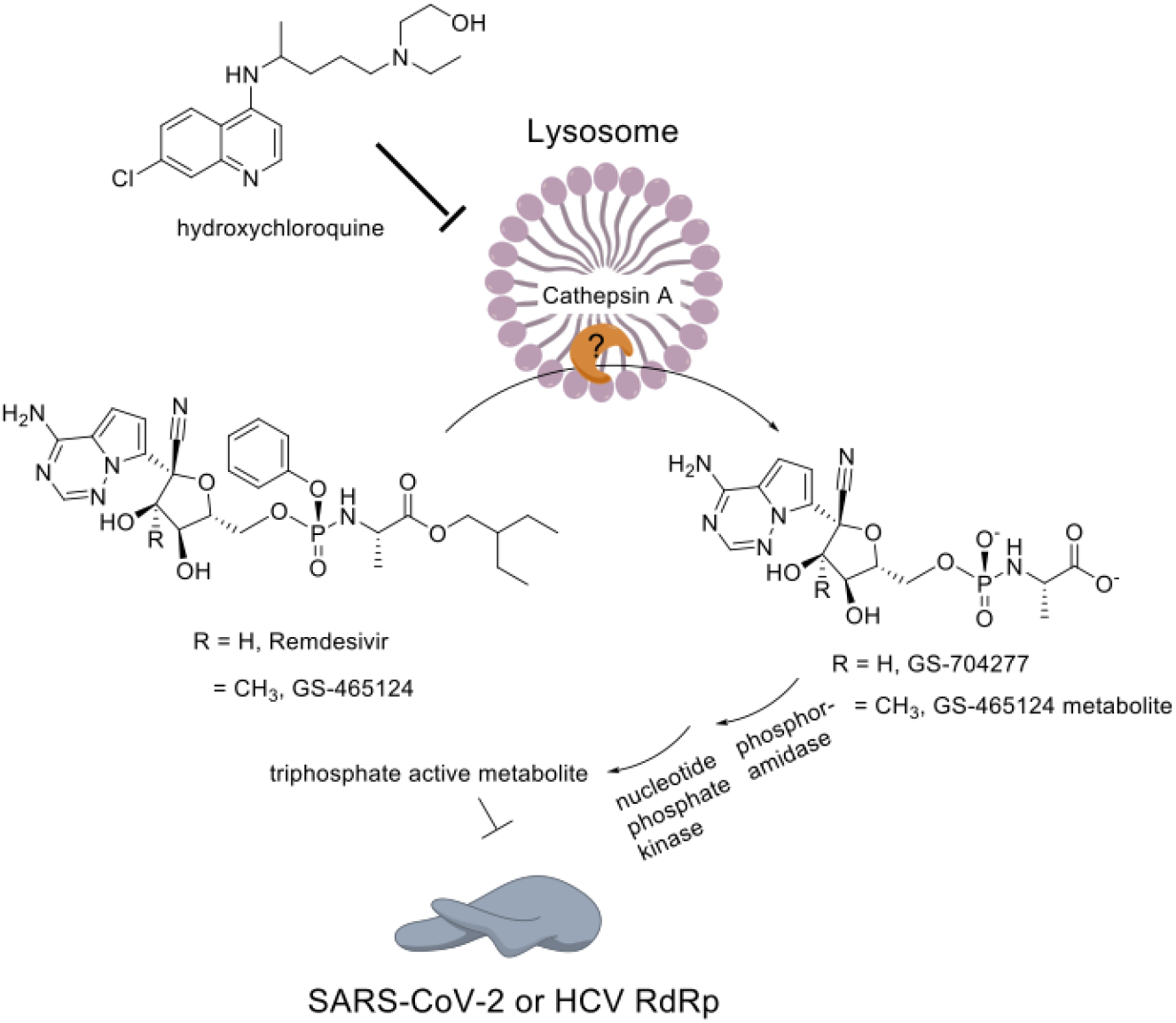
The putative model explaining the antagonism between remdesivir and lysosomotropic agent.

It has been demonstrated previously^63^ that network-based approaches investigating shared biochemical pathways between possible drug combinations are helpful in predicting synergistic combinations and may be useful in predicting the mechanism of action behind observed drug synergism/antagonism. COVID-KOP, a tool recently developed by our group,^36^ integrates existing biochemical knowledge with that contained in literature on COVID-19 in the form of knowledge graphs. Analysis using COVID-KOP also suggests that the mechanism of action for the antagonism of remdesivir and hydroxychloroquine could be specific to the pathways involved in viral entry/egress from cells, as well as in lysosomal acidification (“pH reduction” term), the latter which is consistent with the activation pathway hypothesis detailed above. Additionally, these two drugs were also associated with protein kinase activity, and specifically negative regulation of kinase activity, indicating that a protein kinase recruited during viral infection could interfere with metabolic products of either drug, resulting in antagonism. A brief literature search showed that hydroxychloroquine is an established inhibitor of glycogen synthase kinase-3β (GSK-3β),^70–72^ a serine-threonine kinase that is known to regulate the replication of viruses including dengue virus-2^73^ and SARS-CoV.^70^ In a recent study, in the later stages of infection with dengue virus-2, viral titres were shown to be reduced upon inhibition of GSK-3β.^73^ It has likewise been demonstrated by Wu et al.^70^ that GSK-3 regulates the lifecycle of SARS-CoV and is important in SARS-CoV N protein phosphorylation. Wu et al.^70^ further demonstrated that treatment with kenpaullone, a GSK-3β inhibitor, in turn downregulated SARS-CoV RNA synthesis, and thus hypothesized that phosphorylated N protein likely constitutes part of the viral replication complex in coronaviruses.^74,75^ While remdesivir has no pertinent links to GSK-3β in the literature, it is known that remdesivir inhibits the action of the SARS-CoV-2 RNA-dependent RNA polymerase (RdRp).^76^ Thus, our computational analysis suggests that hydroxychloroquine’s inhibition of GSK-3β may play a role in its antagonism of remdesivir.

It should be mentioned that these biological process and activity terms do not provide further information on what specific protein kinase is involved. Thus, we suggest that these stages of the viral lifecycle, as well as kinase activity during SARS-CoV-2 infection, should be investigated in further explorations of antagonism mechanisms for hydroxychloroquine and remdesivir.

Another compound tested in combinations in our study was amodiaquine, a potent 4-aminoquinoline compound clinically used for the treatment of malaria. It is structurally related to chloroquine, and both are suspected to act through the blocking heme detoxification in the parasite digestive food vacuole (a lysosome like, acidic compartment central to the metabolism of the parasite where hemoglobin is degraded to provide amino acids for parasite metabolism^77^). Indeed, amodiaquine has been reported to interact with μ-oxo dimers of heme ((FeIII-protoporphyrin IX)_2_O) in vitro. The major active metabolite of amodiaquine, monodesethyl-amodiaquine, is rapidly produced via hepatic P450 enzyme conversion in vivo. This metabolite has a half-life in blood plasma of 9–18 days and reaches a peak concentration of ~500 nM 2 hours after oral administration. By contrast, amodiaquine has a half-life of ~3 hours, attaining a peak concentration of ~25 nM within 30 minutes of oral administration.^78^

We have observed a notable synergism for combinations of nitazoxanide with remdesivir, umifenovir, or amodiaquine (Fig. 9). Nitazoxanide is an FDA approved, bioavailable, broad-spectrum anti-infective drug, which recently has been investigated for use against SARS-CoV-2 owing to its previously established anti-coronaviral activities.^79^ It was originally identified as a potential antiviral drug repurposing candidate against SARS-CoV-2 with an IC_50_ of about 2 μM in a focused compound screening including remdesivir and chloroquine.^76^ Previous studies have shown that nitazoxanide is able to activate PKR or/and RIG-I, thus inducing type-I interferon production and signalling in Ebola and vesicular stomatitis virus (VSV).^80^ SARS-CoV-2 is known for its unique immunopathology of reduced production of, but extra sensitivity to, type-I/III interferon.^81^ Interferon-β1b combined with antiviral cocktail (lopinavir-ritonavir and ribavirin) has shown promising synergy in shortening the duration of viral shedding according to a Phase II randomized trial for early COVID-19 treatment.^26^ Taken together, current knowledge suggests an interferon-inducing mechanism of nitazoxanide in vitro. This putative mechanism is also consistent with the broad-spectrum synergy in our matrix screen.

The concentration at which we observed synergy with remdesivir (>1.25 μM, equivalent to 0.383 mg/L) is achievable in plasma and lung trough even at low doses.^82^ Nitazoxanide is well tolerated and there is no report of any significant adverse effects from healthy adults.^83^ As of June 18, 2020, there are 18 nitazoxanide trials (used as monotherapy or combination with other antivirals) registered in clinicaltrials.gov for COVID-19. Moreover, the combination of nitazoxanide with remdesivir looks the most promising from clinical perspective because both drugs would potentially be available for use for the treatment of COVID-19 (nitazoxanide is FDA-approved and remdesivir has an emergency use authorization).

Several drug combinations described in this study are currently being evaluated in clinical trials. The combinations hydroxychloroquine-favipiravir^84–86^ and hydroxychloroquine-lopinavir/ritonavir^87^ were reported. However, all the trials above are still recruiting or not yet recruiting. Given the common use of some of our compounds in clinical trials, it might be possible that more are being tested in clinical trials as part of standard of care (SOC) inclusion. This means that these drugs are being tested for their efficacy in tandem with the SOC, which could contain hydroxychloroquine, or remdesivir, or other compounds, depending on the time and place.

Despite strong synergism and antagonism demonstrated by drug combinations reported in this study, we would like to emphasize that these results require further validation. Vero E6 cell do not express the serine proteases TMPRSS2/4, the two proteases crucial for viral entry through the early membrane fusion pathway of invasion (e.g., nafamostat, camostat, and lysosomotropic agents).^88^ Thus, all synergistic and antagonistic combinations need to be verified in other cell lines to determine how penetrant the reported synergies are, for example, in a TMPRSS2/4 expressing cell line or primary airway cell model. The synergy between amodiaquine and nitazoxanide in Vero E6 may not translate to Calu-3 (TMPRSS2^+^) where single-agent amodiaquine is >10-fold less potent.^88^ Moreover, the observed synergism/antagonism may not be maintained *in vivo* due to complex pharmacokinetics; meanwhile, antagonism sometimes can be circumvented by altering dosing schedule or formulation. Therefore, more in-depth *in vivo* validation and pharmacokinetic modelling are still necessary.

Here, we observed a common antagonism between lysosomotropic amines and remdesivir, although some drugs (such as umifenovir, mefloquine, and amodiaquine) appeared to synergize with remdesivir at high concentrations. This synergy at high concentrations may not translate well to a clinical setting, given (1) lysosomotropic agents have been shown ×10 less potent in TMPRSS2/4^+^ cell model;^88^ (2) it is challenging to maintain a high steady-state plasma concentration without introducing additional toxicity; and (3) the concentration at which shows synergy may not be achievable *in vivo*. For example, 10 μM umifenovir is unlikely achievable in humans (Cmax = 467 ng/mL, equivalent to 0.98 μM).^89^

Principally, these findings suggest that lysosomotropic amine drugs, such as hydroxychloroquine, should be prescribed and used with increased caution in COVID-19 patients due to their relatively long half-life (usually weeks). This is consistent with a statement issued by the FDA on June 15, 2020, warning that combinations of chloroquine/hydroxychloroquine with remdesivir may reduce the antiviral effectiveness of remdesivir against SARS-CoV-2,^90^ based on an unnamed, independent, *in vitro* experiment.

We want to emphasize, that in addition to 73 binary combinations described in this paper, we also identified in silico 95 ternary combinations of 15 drugs.^29^ However, experiments if ternary combinations are more complex, thus, testing those we are considering as a future study.

## Conclusions

Using *in silico* approaches and our expertise in data science, cheminformatics, and computational antiviral research, we recently identified 281 combinations of 38 drugs with potential activity against SARS-CoV-2 and prioritized 73 combinations of 32 drugs as potential combination therapies for COVID-19. Experimental evaluation of these combinations via cytopathic effect and host cell toxicity counter assays in 6×6 dose matrix allowed us to identify 16 synergistic and 8 antagonistic combinations, with four of them exhibiting both synergy and antagonism.

Despite strong synergism/antagonism observed in some cases here, these are from cell culture experiments thus requiring more in-depth validation. Among 16 synergistic cases, nitazoxanide combined with three other compounds (remdesivir, amodiaquine, and umifenovir) exhibited significant synergy against SARS-CoV-2. Notable synergism was also demonstrated by combining umifenovir with mefloquine, amodiaquine, emetine, or lopinavir. Nitazoxanide – remdesivir combo looks the most promising from clinical perspective because both drugs are approved by FDA (remdesivir through Emergency Use Authorization).

Among eight observed cases of antagonistic interaction, the most notable is the strong antagonism demonstrated by combination of remdesivir and hydroxychloroquine, which is currently undergoing clinical trials. In four cases, i.e., remdesivir + mefloquine, remdesivir + amodiaquine, remdesivir + umifenovir, and nitazoxanide + lopinavir, both antagonism and synergism were observed; the effect was concentration dependent.

Altogether, our findings demonstrate the utility of *in silico* tools for rational selection of drug combinations, the importance of preclinical testing of drug combinations prior to their administration in patients, and the overall promise of using drug repurposing and combination therapies against SARS-CoV-2.

All the protocols and results are freely available to scientific community at https://opendata.ncats.nih.gov/covid19/matrix.

## Acknowledgements

Data-mining tools used in this study were developed under the Biomedical Data Translator Initiative of National Center for Advancing Translational Sciences, National Institute of Health (grants OT3TR002020, OT2R002514) and under support of National Institute of Health (grant 1U01CA207160). This research was also supported in by the Intramural Research Programs of the National Center for Advancing Translational Sciences (NCATS), National Institutes of Health (NIH).

